# Motor Learning Mechanisms are not modified by Feedback Manipulations in a Real-World Task

**DOI:** 10.1101/2024.04.10.588812

**Authors:** Federico Nardi, A. Aldo Faisal, Shlomi Haar

## Abstract

Error-based and reward-based mechanisms of motor learning co-occur in real-world scenarios but are traditionally isolated in laboratory tasks via feedback manipulations. This study examines the distinctiveness of these mechanisms by applying a lab-based feedback manipulation to a real-world task. Using Embodied Virtual Reality (EVR) of pool billiards - allowing for full proprioception via interaction with the physical pool table, cue stick, and balls - we introduced visual perturbations to a real-world task. 32 participants (12 F) underwent two sessions learning a visuomotor rotation, once with error and once with reward feedback. While naive participants corrected the entire rotation with error feedback, only partial correction was observed with reward feedback, highlighting the influence of the feedback regime on learning. However, the reward-dependent motor variability, lag-1 autocorrelation decay, and inter-trial variability decay - all indicators of reward-based and skill learning - were higher in the error feedback session, suggesting that the provided visual feedback did not exclusively engage specific learning mechanisms. Analysis of post-movement beta rebound (PMBR), a brain activity marker of learning mechanisms, revealed a decrease in PMBR with reward feedback but no consistent trend during error feedback sessions. These findings support the behavioural results, suggesting that while reward feedback was absent in error conditions, participants still engaged in reward-based learning. This study underscores the complexity of motor learning processes and highlights that visual feedback by itself can not elucidate the interplay between error-based and reward-based mechanisms in real-world contexts.

## 1 Introduction

Motor learning plays a crucial role throughout our lifespan, shaping human behavior from a baby learning to walk to a person living with Parkinson’s disease adapting their movements to compensate for tremors.

This learning process is considered to be driven by two key mechanisms: *error-based* learning, which relies on quantifiable sensory-prediction errors, and *reward-based* learning, which reinforces successful actions Wolpert et al. (2011). For instance, when playing billiards, we use *error-based* learning to correct our next shot based on the disparity between the pocket and the actual outcome. Conversely, after pocketing the ball, *reward-based* learning positively reinforces the successful actions, increasing the likelihood of repeating the same movements in the future.

This dichotomy in motor learning mechanisms has been extensively studied in the field of neuroscience. Specifically, *error-based* learning is considered to be based on adaptation mechanisms in the cerebellum Kawato and Gomi (1992); Doya (2000); Taylor and Ivry (2014), while *reward-based* learning is often associated with the brain’s reward pathways in the basal ganglia and the release of neurotransmitters such as dopamine, which play a crucial role in encoding the value of particular motor actions Dayan and Balleine (2002); Niv (2009); Diedrichsen et al. (2010).

Understanding the interplay and influence of these error-based and reward-based learning mechanisms is crucial for developing effective strategies to enhance motor learning and skill acquisition Krakauer (2006). Exploring their interaction in realistic and diverse settings can provide a deeper understanding of how individuals learn and adapt their movements in everyday life Wolpert et al. (2011). Typically, these two processes of error-based and reward-based learning are studied as distinct mechanisms in simple laboratory tasks that allow for isolating them through controlled feedback manipulations Taylor and Ivry (2014); Therrien et al. (2016). However, the unparalleled complexity of human movement cannot be fully captured by such simplistic tasks. Our recent work has shown that in a real-world task of playing billiards, error-based and reward-based learning coexist, with participants using a combination of both mechanisms while predominantly relying on one or the other Haar and Faisal (2020).

Recent advancements in real-world neuroscience have enabled the study of neurobehavioral processes in natural human behavior, offering new insights into the complexities of motor learning and skill acquisition Haar et al. (2020); Stout et al. (2021); Krotov et al. (2022); Shamay-Tsoory and Mendelsohn (2019). However, these approaches come with limitations, as controlling different types of feedback and learning mechanisms is more challenging compared to classical lab-based tasks. To overcome this limitation, we integrated our billiards task into an Embodied Virtual Reality setup. This setup allows for visual manipulations without compromising the complex movements required by a real-world task Haar et al. (2021). The EVR setup synchronises a physical pool table, cue stick, and balls with a virtual environment using optical marker-based motion capture technology. Participants can interact with physical objects simultaneously in both the real and virtual worlds, enabling us to manipulate the visual environment while they engage in realistic movements accompanied by full proprioceptive feedback Nardi et al. (2023).

In this study, we leveraged our Embodied Virtual Reality setup to explore the potential of feedback manipulations commonly used in lab-based tasks to differentiate error-based and reward-based learning mechanisms in a real-world motor learning task. We hypothesised that by eliminating specific visual information in the EVR setting, participants would exclusively rely on the relevant learning mechanism, resulting in observable differences in their behavior and neural patterns during the training sessions. Our analysis of movement data and adjustments in brain activity following these feedback manipulations provides new insights into the connection between visual feedback and the distinct characteristics of error-based and reward-based learning mechanisms in the context of real-world human behavior.

## 2 Methods

### 2.1 Experimental Setup

Our experimental setup combines a physical pool table with Embodied Virtual Reality to accurately measure and adjust participants’ actions. Participants play an actual pool game while wearing an HTC Vive headset and receive customised visual feedback in the virtual environment, tailored to the targeted learning mechanism (fig. 1a). The physical cue stick used in the game is faithfully replicated in VR, with its geometric characteristics and marker positions streamed through high-precision Optitrack cameras (accuracy of *±*0.2mm), enabling participants to perceive it as a virtual object.

**Fig 1.**
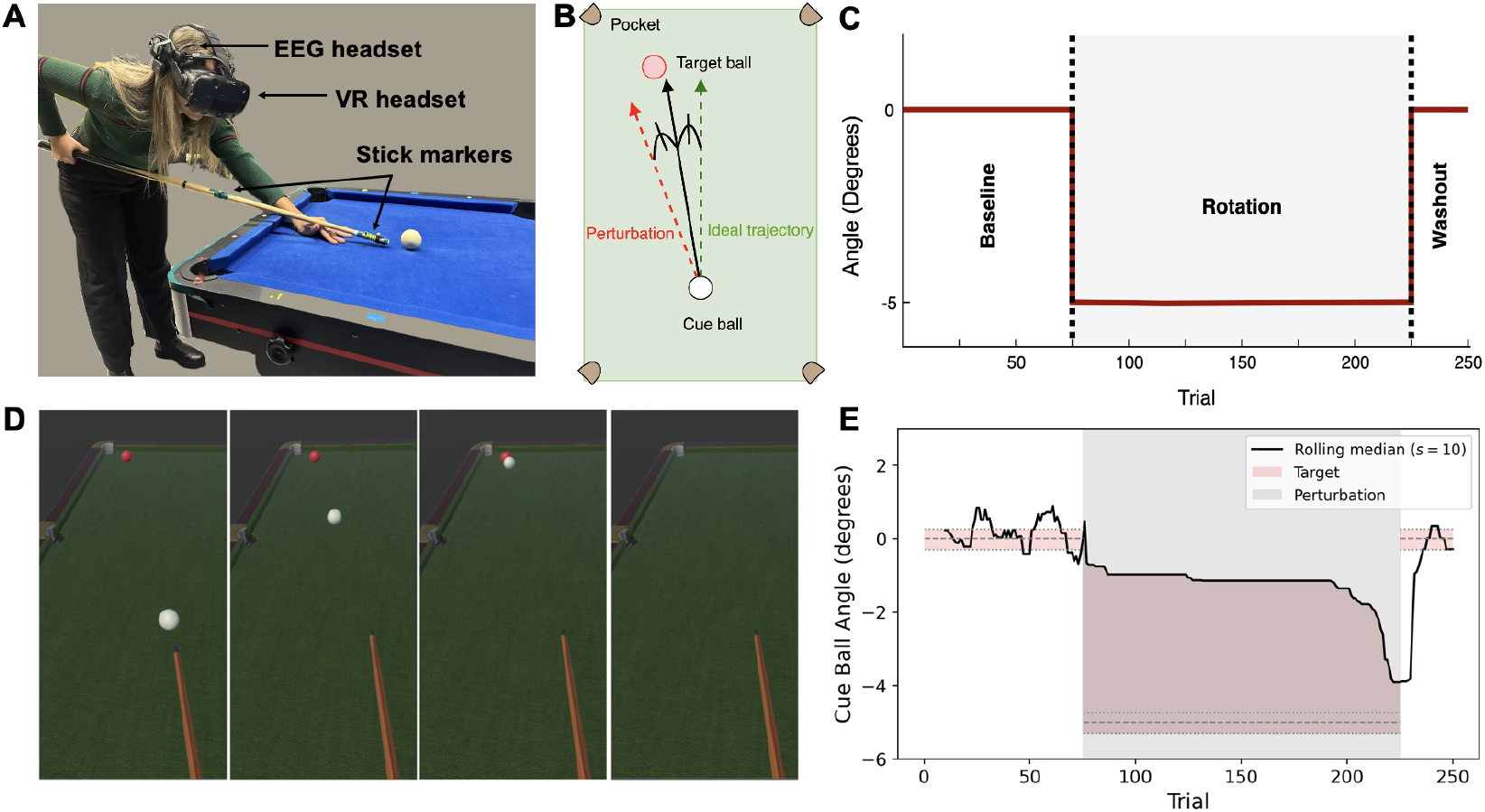
Experimental Paradigm. **(A) Participant playing in the EVR setup. (B) Perturbation** applied during the *Rotation* phase: the trajectory of the cue ball is rotated 5° toward the side of the table. **Structure of the experiment**: after 75 trials of full feedback and no perturbation (*Baseline*), a visuomotor rotation is applied on the cue ball trajectory (*Rotation*) for 150 trials; follows a final phase of 25 trials without perturbation and full feedback (*Washout* ). The grey area represents the presence of perturbation, whereas the solid red line represents the ideal angle. **(D) Visual feedback for the error task**. At collision, the cue ball and the target ball disappear to make the participant correct only based on the trajectory before the hit. **(E) Reward zone** of an example participant. The grey area represents the presence of perturbation, while the solid black line represents the median of the last 10 rewarded trials. The pink area represents the zone that rewards the participant when the shot is directed in any part of the area: during *baseline* and *washout* it corresponds to the pocketing funnel, while during *rotation* the reward zone is inflated to direct the participant towards the new ideal trajectory. The dotted lines show the range of successful angles, in all the phases of the experiment.

This embodiment allows participants to physically interact with real-world objects, experiencing complete somatosensory and proprioceptive feedback. Participants shoot a physical cue ball, maintaining haptic feedback and realistic collision sounds. However, artificial sounds represent collisions between the cue ball and target balls, which are solely virtual but programmed to adhere to real physics like other objects in the virtual environment (for validation see Haar et al. (2021)). Pre-experiment calibration using four cameras and VR controllers ensures alignment between the dimensions of the virtual and real tables, enhancing the realism.

The Embodied Virtual Reality paradigm was developed using Unity software programmed in C#. Additionally, the setup integrates a 14-channel wireless electroencephalogram device (eMotiv EPOC+) to record brain activity during the experiments. This device can be easily worn alongside the VR headset without interference.

### 2.2 Experimental Design

332 healthy volunteers (20 men and 12 women) aged 20 to 25 with normal or corrected-to-normal vision and little to no experience in playing pool, participated in an experiment. The experiment included two sessions which took place on different days, 24 to 48 hours apart. Each session (fig. 1c) consisted of a *Baseline* phase (75 trials, divided into 3 blocks of 25 trials), a *Rotation* phase (150 trials, comprised of 6 blocks of 25 trials) and a final *Washout* block (25 trials). In each trial, participants took a single shot, aiming to pocket a target ball placed near the far-left or far-right corner of the pool table, starting from a fixed position. Without any time pressure, the trials ranged in duration from several seconds to over ten seconds, depending on the individual’s pacing.

During the *Rotation* phase, a 5° visuomotor perturbation was applied to the cue ball’s trajectory (Fig. 1b), compelling participants to adjust their aim toward the table’s centre to successfully pocket it. Specific feedback related to the condition was provided through virtual reality during this rotation phase. The task was structured to encourage participants to rely on a particular learning method by concealing visual information that contributed to the opposite mechanism.

In the error-based feedback condition, the ball trajectories were hidden after colliding with each other, forcing participants to rely solely on cue ball trajectory errors for progression, as they could not see the target ball being pocketed. Conversely, in the reward feedback condition, the balls were hidden after the cue ball was shot. Successful shots resulted in participants being shown an artificially generated successful trajectory as a form of reinforcement, according to well-established motor learning principles Therrien et al. (2016, 2018); van Mastrigt et al. (2023).

The reward region was established using two specific criteria: (i) shooting within a successful zone around the pocket, or (ii) when the accuracy of the shot exceeded the median of the previous 10 successful attempts. The defined successful trials assisted participants in making necessary adjustments without error feedback.

A previous pilot study aimed to determine the optimal level of reinforcement. Providing too many rewards would allow participants to quickly reach the correct path, but may reduce their interest. Conversely, setting the reward threshold too high would make it difficult for participants to adjust in a reasonable time due to a lack of guidance. To verify this, several participants took part in pilot experiments where trials were considered “successful” if they exceeded a specific percentage of earlier rewarded trials. The optimum threshold was found by surpassing both the 40^*th*^ percentile and median performance levels, which were close; therefore, we prioritised meeting the median requirement for easier explanation without significant compromise.

### Behavioural Data Analysis

The data were pre-processed to address extreme values. Specifically, a threshold was established to remove outliers in terms of cue ball directional errors during the baseline (*µ* ± 3*σ*), and it was applied as well in the perturbation learning zone.

To enhance the robustness and reliability of detected trends, all primary results were evaluated by grouping trials into blocks of 25. Additionally, no significant difference was observed between target pockets, enabling the aggregation of samples (aiming to the left or right pockets) for increased statistical power. However, some differences emerged between participants performing the task in the first or second session with the same feedback. Therefore, when appropriate, the groups were separated between sessions for analysis.

The primary focus of the behavioural data was on the directional error made by the participants in each shot. After defining the ideal successful trajectory (set as 0°), the angle of the cue ball was measured by centering the range around that value and analysing the deviations in the direction of the shot.

The trial-by-trial directional error was used to measure learning. When the visuomotor rotation was introduced during the rotation phase, the “ideal” angle changed from 0° to -5° due to the rotation, and all results are presented accordingly (fig.1c). To determine the best fit between linear and double exponential for the learning curves of the two feedback conditions, we performed an F-test comparing the residual sum of squares of the two curves. The parameters of the curves are reported in the results and referred to as *τ*_fast_ and *τ*_slow_, respectively, for the fast and slow components of the exponential curve. The linear fit is reported through *b*_0_ and *b*_1_, representing respectively the intercept and slope of the linear model.

Another aspect of the game that was considered was the success rate, which had two distinct components: the *real* success rate and the *perceived* success rate. During the baseline and washout phases, when no changes were introduced, these two rates were aligned. In the perturbation phase of the error feedback task, they were aligned as well because participants lacked visual information about the success of their shots. Conversely, in the reward feedback session, the two rates differed significantly, as the perceived success was based on the rewards given according to the reward zone, while the real success referred to the actual instances when the target ball was pocketed.

To account for uncertainty in the shots and maintain consistency with previous research, the variability of the shots was analysed. Specifically, the focus was on measuring the corrected variability, which involved calculating the standard deviation of the residuals after removing any linear trend from the directional error in shots at a block level. This approach remained valid even when an overall exponential trend was present at the session level, as a sufficiently linear trend was observed within blocks of 25 trials.

The methodology involved the use of mixed-effects ANOVA and mixed-effect linear models to analyse the repeated measurements from each participant. Non-parametric tests, such as paired Wilcoxon Signed-Rank tests and Mann-Whitney tests, were utilised for sample testing and comparisons to account for non-normal distributions of the samples. Specifically, Wilcoxon Signed-Rank tests were employed to compare single samples against a 0 median or paired samples, while Mann-Whitney tests were used to compare the different samples.

The mixed effects ANOVA tests reported in the results sections refer to the baseline (*F*_•|base_), perturbation (*F*_•|pert_), or washout phases (*F*_•|wash_), and were executed between different modes (*F*_mode|•_) or the interaction between mode and time (*F*_int|•_).

### Behavioural Measures of Learning

To further validate the behavioural results obtained with the two individual mechanisms, three wellknown measures of motor learning were used to demonstrate differences between the feedback: lag-1 autocorrelation, inter-trial variability, and reward-dependent motor variability.

The lag-1 autocorrelation (ACF) is calculated by correlating the directional error time series of one session with itself, with a one-time step lag. To address potential bias in autocorrelation estimation from short time series Arnau and Bono (2001), we analyse autocorrelation functions for sets of trials during the learning session at different stages: baseline (blocks 1-3), early learning (blocks 4-6), and late learning (blocks 7-9) rather than within each set of 25 trials.

Inter-trial variability (ITV) was assessed at the block level, considering the corrected variability across the 25 trials within each block. We assessed the decay of ITV by comparing the baseline (last block trials 51 to 75), early learning (first block 76 to 100), and late learning phases (last block - 201 to 225).

The reward-dependent motor variability is derived by comparing the trial-to-trial changes in performance after a success or a failure. This measure is traditionally linked to reinforcement-based motor learning and has been shown to increase after unrewarded trials, reflecting the system’s effort to explore new motor strategies to find success Pekny et al. (2015). The reward-dependent motor variability was calculated considering the distributions of changes in shot angle after successful and unsuccessful trials for each participant.

### EEG Acquisition and Pre-processing

The EEG data was collected using an Emotiv EPOC+ headset, a wireless 14-channel system positioned under the VR headset. The data was sampled at 256Hz and aligned with the VR data through the trial timestamps. The entire pre-processing pipeline was executed in MATLAB using the EEGlab Delorme and Makeig (2004) and Fieldtrip Oostenveld et al. (2011) toolboxes.

After applying high-pass and low-pass filtering with a FIR filter at 1Hz and 45Hz respectively, the data was cleaned using the EEGlab *clean rawdata* function. This function automatically removes artifacts from the signals, including bad portions of data and problematic channels, using the *Artifact Subspace Reconstruction* algorithm Kothe (2013). Specifically, it identifies channels with a large amount of noise based on their standard deviation and poorly correlated channels. To retain as much informative data as possible, the default rejection threshold for channel correlation was gradually decreased by steps of 0.05 until no more than three channels were removed or the minimum correlation reached 0.6. This approach aims to minimise the loss of informative data while preventing excessive removal of channels or retention of non-informative data.

Furthermore, the signal windows were removed based on thresholds for the maximum standard deviation of bursts and the maximum fraction of contaminated channels permitted in the final output data for each considered window. The specific parameters were selected through a manual examination of the sessions’ time-frequency plots, evaluating 4 combinations of parameter settings with increasingly strict criteria.

We extracted epochs from the EEG data, focusing on the shots as a valid reference for the participants’ movements, consistent with previous *Event-Related Potential* (ERP) and event-related spectral perturbation (ERSP) evaluations, as well as the work of Haar and Faisal (2020). The epochs were set to 9 seconds in duration around each shot, establishing a baseline reference before and after the movement and including larger temporal margins to accurately detect fluctuations in beta activity. However, since post-movement beta rebound can be detected up to 1000 ms after movement onset Jurkiewicz et al. (2006), we focused only on that interval to identify the peak in beta activity.

We then performed an *Independent Component Analysis* on the data to remove muscle-, eye-, and heart-induced artifacts. Afterwards, we interpolated the C3 channel from the other existing channels using spherical splines and a topological map consistent with the international 10-20 EEG system, which positions C3 over the left motor cortex Jasper (1958). This channel placement was informed by previous studies on motor learning Tan et al. (2014, 2016) and the right-handedness of all participants.

### EEG Time-Frequency Analysis

After preprocessing the EEG data, we conducted a time-frequency analysis to investigate the brain activity associated with the participants’ movements during the experiment, while providing individual feedback. The time-frequency domain transformation was performed on each block using a convolution with complex Morlet wavelets in 0.8 Hz steps and 3 millisecond cycles.

To ensure comparability between various sessions within and between participants, we standardised the data so that all sessions had equal total power in the specific frequency range. Specifically, we calculated the total power within the 8-45Hz range to determine the normalisation factors for each separate session Liu et al. (2021). This normalisation method guarantees that variations in beta are more noticeable and accurate, allowing for precise detection of fluctuations that are solely due to the signal itself, as potential trends and patterns would be observable across all frequencies and not just in the beta band. Furthermore, normalising across sessions eliminates any trends in the baseline, which we define as the median power over each session, as the correction is also applied between the different blocks of the same session.

The subsequent analysis involved calculating the EEG power change in response to events as a percentage change, normalised relative to the median power evaluated in 1Hz steps over each block. This approach was necessary due to the absence of a clear baseline during the task Torrecillos et al. (2015); Tan et al. (2014, 2016). The process involved subtracting one from the normalised value and then multiplying by 100 to obtain a comparable percentage scale measure. To ensure robustness, we calculated the peak value on the average signal across the beta-band frequencies and derived one value per trial. Then, we took the median of these peak values across various trials to obtain a PMBR value for each block.

Additionally, to ensure data reliability, we established criteria to include or exclude specific sessions in the group analysis. EEG data can have missing values, possibly due to electrode misplacement as the task required significant movements. Blocks of an individual session were included in the EEG time-frequency analysis only if at least half of the block (13 trials) had available data. Entire sessions were removed if at least 3 blocks were missing, to accurately capture trends and avoid including noisy information.

After selecting reliable sessions, missing data were imputed separately for error and reward sessions, using *Probabilistic Principal Component Analysis* (PPCA). This method samples from the conditional distribution of missing values given the observed data Tipping and Bishop (1999). The overall data preprocessing reduced the number of participants in the EEG analysis from 32 to 28 and 30, respectively for the error and reward tasks. Finally, all block values were aggregated by block and reported by condition.

## Results

### Learning and success are strongly modulated by visual feedback manipulations

Participants were able to successfully learn and correct for some of the rotation under both error and reward conditions. However, their learning patterns varied significantly due to the feedback manipulation in the EVR setup (see fig. 2A), confirming our preliminary results Nardi et al. (2023).

**Fig 2.**
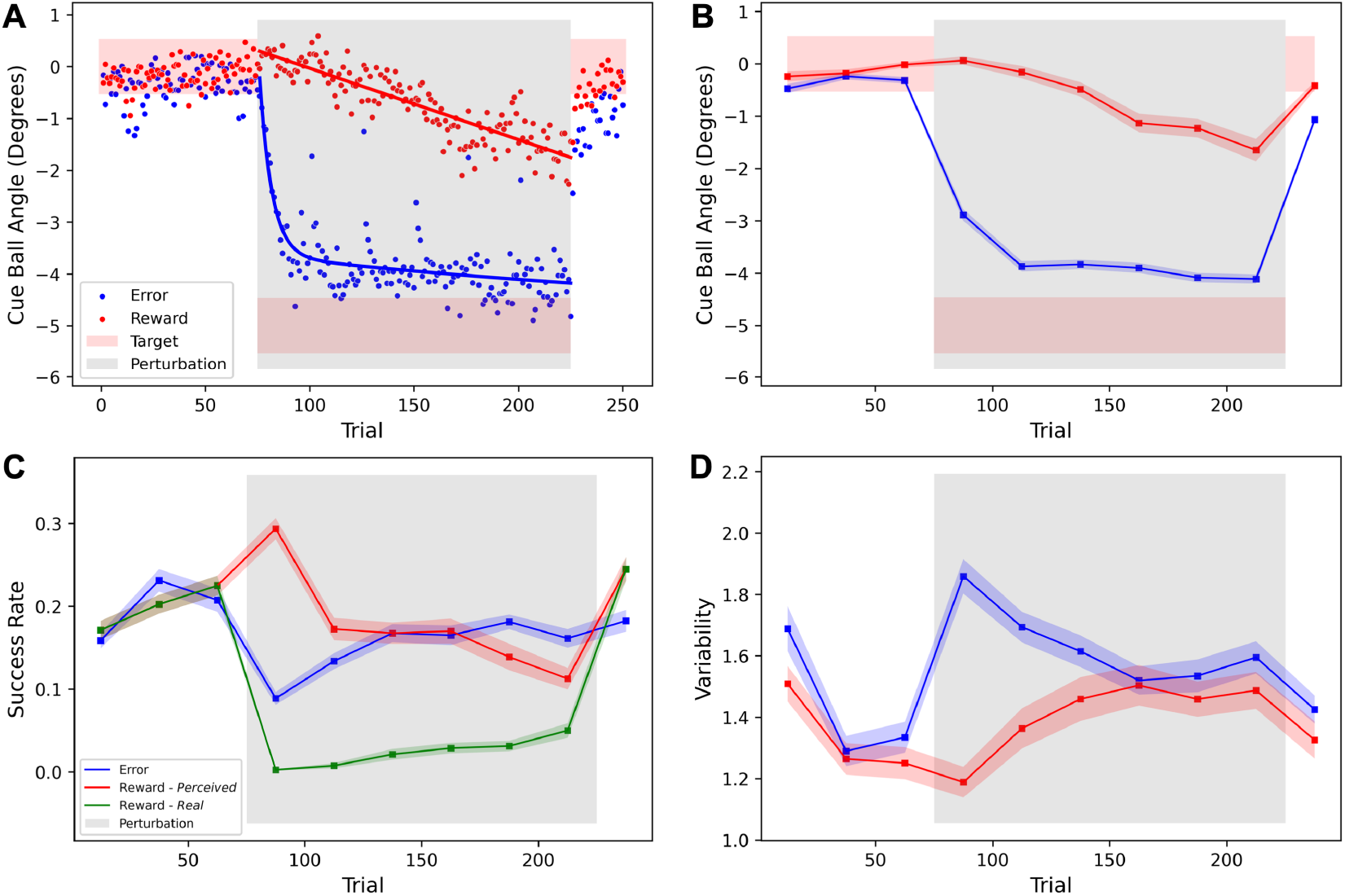
Task performance in error (blue) and reward (red) conditions. **(A)** Average cue ball trial-by-trial angle. The pink area indicates the range of successful angles. The bold line is a double exponential for the error data /linear fit for the reward. **(B)** Average cue ball angle over blocks. **(C)** Success rate for the *error* condition, vs Perceived (red) and Real (green) success rates for the *reward* condition. Corrected variability, after removal of linear trends. **(B-D)** Each block of 25 trials is represented by its distribution, the means over participants are connected by the bold lines and the shaded areas represent inter-participant variability (standard error of the mean). Grey area represents the presence of perturbation. *N* = 32.

During the baseline phase, with full visual feedback and no perturbation, participants exhibited similar performance in both sessions. However, during the rotation phase with partial visual feedback and a perturbation on the cue ball’s trajectory, participants learned the rotation more quickly with error-only feedback. This resulted in a larger after-effect during the washout phase. The trial-by-trial directional errors in the error-feedback session showed a fast initial learning phase followed by a sustained slower one, best captured by a double exponential curve (*τ*_fast|error_ = 6, *τ*_slow|error_ = 296). In contrast, learning through reward-only feedback progressed at a slower pace, with a linear curve providing the best fit (*b*_0_ = 0.31, *b*_1_ = −0.014).

The block averages displayed a similar pattern (see fig. 2B), with a significant difference between the two learning trends during the perturbation phase (*F*_mode|pert_(1, 62) = 96.25, *p <* 0.001; *F*_int|pert_(5, 310) = 3.94, *p <* 0.01). Despite learning at a slower rate in the reward condition, participants were still able to compensate for at least some of the visuomotor rotation in both conditions.

The slower learning with reward-only feedback can be partially explained by the use of a “reward zone,” which led to an increased perceived success in the early blocks of the rotation. In an error feedback session, a successful shot is simply defined as when the trajectory was heading towards pocketing. With reward-only feedback, there are two types of success: *real* and *perceived*. While real success is equivalent to that in the error feedback condition, where the trajectory was heading towards pocketing, perceived success is based on the received rewards. Early on in rotations, rewards are given whenever the direction is closer to reaching for a rotated pocket than for the original one.

Accordingly, success rates were similar between sessions in the baseline and washout blocks (*F*_mode|base_(1, 62) *<* 0.01, *p* = 0.99; *F*_mode|wash_(1, 62) = 2.57, *p* = 0.12). However, during the perturbation phase we observed very different success profiles (fig. 2C). The real and perceived success rates during reward-only blocks showed opposite trends (*F*_int|pert_(5, 310) = 7.54, *p <* 0.001), and the success rates during error-only blocks differed from both perceived (*F*_int|pert_(5, 310) = 10.11, *p <* 0.001) and real reward (*F*_mode|pert_(1, 62) = 52.33, *p <* 0.001).

The variability in shot direction under different conditions (fig. 2D) closely corresponded with their perceived success rates. To address potential learning trends within a block, we calculated the *corrected variability* by determining the standard deviation of residuals from a block-level linear regression model applied to the cue ball angle dataHaar et al. (2020). The participants’ performance exhibited similar patterns during the baseline and washout phases (*F*_mode|base_(1, 62) = 0.43, *p* = 0.52; *F*_mode|wash_(1, 62) = 0.43, *p* = 0.52), but demonstrating opposite dynamics during the perturbation phase over time (*F*_int|pert_(5, 310) = 6.90, *p <* 0.001).

Comparing the different sessions within the individual feedback conditions, we observed distinct differences across sessions in the reward task. The average cue ball angle over time (Fig.3B) shows a clear distinction between sessions. The participants in Round 1 - who were naive to the task and the EVR - did not learn how to correct for the perturbation, whereas participants in Round 2 - who were less naive to the task and the EVR as they already had a 1st session where they aimed at the other pocket with error-feedback - corrected for most of the perturbation (2.82 *±* 0.29 degrees), having results not significantly different from both sessions of the error task (*W*_*ErrR*1_ = 34, *p* = 0.083; *W*_*ErrR*2_ = 33, *p* = 0.074). The same was found for the success rate (Fig.3C), which aligns with the learning of the task. Participants who performed the task in their first session rarely experienced real successes once the perturbation was introduced, while those who performed it in their second session (thus, were not completely naive) improved their real success rates over learning, reaching up to 10% of their shots.

**Fig 3.**
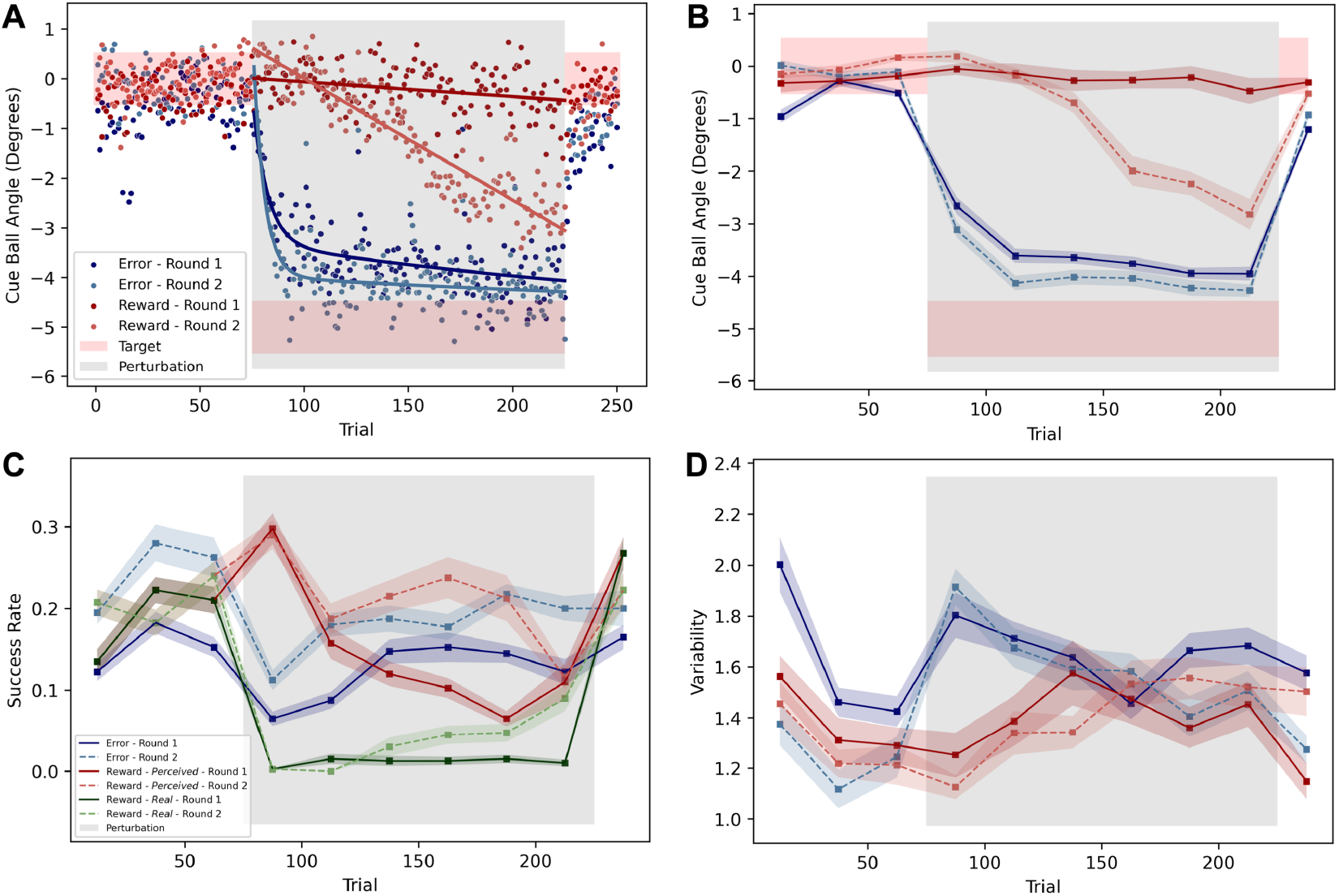
Task performance by sessions: First (dark and solid) vs Second (light and dashed) in error (blue) and reward (red) conditions. **(A)** Average cue ball trial-by-trial angle. The pink area indicates the range of successful angles. The bold line is a double exponential for the error data /linear fit for the reward. Average cue ball angle over blocks. **(C)** Success rate for the *error* condition, vs Perceived (red) and Real (green) success rates for the *reward* condition. **(D)** Corrected variability, after removal of linear trends. **(B-D)** Each block of 25 trials is represented by its distribution, the means over participants are connected by the bold lines and the shaded areas represent inter-participant variability (standard error of the mean). Grey area represents the presence of perturbation. *N* = 32.

**Fig 4.**
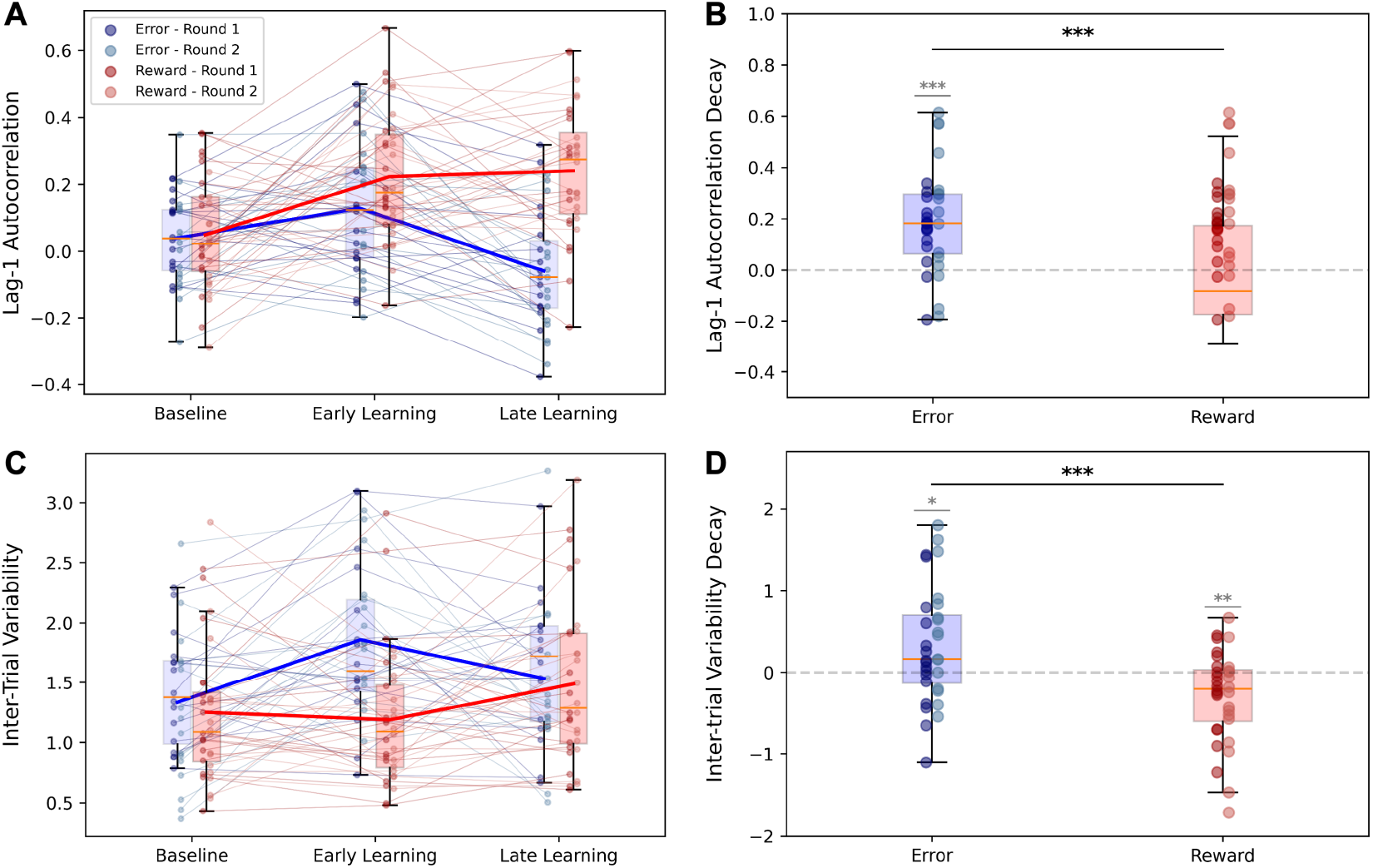
Behavioural differences between error (blue) and reward (red) conditions. **(A)** Lag-1 autocor-relation during *Baseline* (Trials 1-75), *Early Learning* phase (trials 76-150) and *Late Learning* (trials 151-225); **(B)** Decay of lag-1 autocorrelation between *early* and *late* learning, during the *rotation* phase; Inter-trial variability in the last baseline block (*Baseline* - trials 51-75), first block of rotation block (*Early Learning* - trials 76-100) and last rotation blocks (*Late Learning* - trials 201-225); **(D)** Inter-trial variability decay from *Early* and *Late* learning. The darker dots represent *Round 1*, while the lighter *Round 2*, for both error and reward. *N* = 32. **(A-C)** Solid lines represent the averages considering both rounds. **(B-D)** were tested for significant differences against 0 (in grey) and between conditions (in black). ^∗^ : *p <* 0.05, ^∗∗^ : *p <* 0.01, ^∗∗∗^ : *p <* 0.001

The variability of the shots when separating the *First* and *Second* rounds (Fig.3D) showed similar trends between sessions, unlike the other measured quantities.

### Behavioural Metrics suggest mixture in Learning Mechanisms

To assess the contribution of reward-based reinforcement learning and error-based adaptation in the two feedback conditions, we looked at metrics of skill and adaptation learning, which are traditionally linked to reward and error mechanisms, respectively.

We initially analysed the lag-1 autocorrelation [ACF(1)] of the directional error of the cue ball. The decay of autocorrelation is considered a characteristic of skill learning, which is typically associated with reward-based learning rather than error-based adaptation van Beers et al. (2013). To quantify the change in ACF, we examined its decay between the two halves of the perturbation phase. Interestingly, a statistically significant decrease in ACF was found in the error-only task, as tested with a Wilcoxon Signed-Rank test (*W* = 45, *p <* 0.001), indicating skill enhancement even without visible reward feedback. In contrast, there was no significant change in ACF for participants during the reward-only task as learning progressed from early to late stages (*W* = 224, *p* = 0.47).

The pattern observed from baseline (trials 1-75) to early learning (trials 76-150) remained consistent across both sessions, aligning with the perturbation provision and the decline in learning observed in the behavioural results. Furthermore, a significant difference in the pattern was detected between the two feedback conditions, as confirmed by a 2-sample Wilcoxon Signed-Rank test (*W* = 92, *p <* 0.001).

The second metric we investigated was the decay in inter-trial variability, which is known to be a distinctive feature of reward-based skill learning Müller and Sternad (2004); Cohen and Sternad (2009); Shmuelof et al. (2012); Sternad (2018); Krakauer et al. (2019). We quantified this by calculating the difference in corrected variability between the first and last blocks of the perturbation phase, as a measure of the uncertainty throughout the learning period.

Surprisingly, in line with the lag-1 autocorrelation findings, we observed a positive decay in variability when providing error feedback (*W* = 142, *p* = 0.02), while the opposite trend was displayed for the reward feedback task (*W* = 122, *p <* 0.001). We also obtained a significant difference in this metric between the two conditions (*W* = 97, *p* = 0.001), whereas no differences were detected during the final baseline block (*W* = 225, *p* = 0.48).

When examining the reward-dependent motor variability within the error and reward feedback conditions (Fig.5A), we observed significant differences in the changes of shooting angle between trials following success and failure, for both the error and reward tasks across all the measured quantities. As expected, participants made greater corrections after a failed shot rather than after a successful (rewarded) one (Fig.5A first panel), which led to increased exploration when using reinforcement-based learning. This difference was present in both tasks, validating the hypothesis that reward-based learning was also occurring in the error feedback condition.

**Fig 5.**
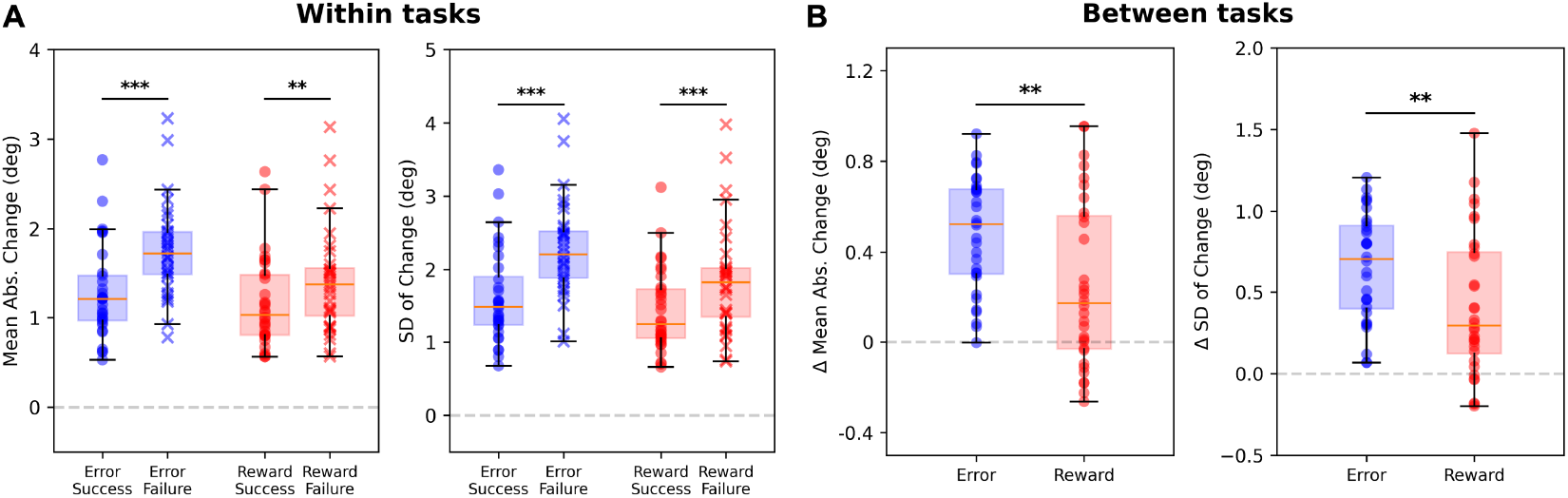
Trial-to-trial angle change after success/failure. **(A)** Comparison of the average quantities for *error* task - after success (blue dots) and failure (blue crosses) - and *reward* task - after success (red dots) and failure (red crosses). **(B)** Difference (Δ) between changes after failures and successes for the *error* and *reward* tasks. The Δ is calculated by subtracting the average change after success and after failure trials, for each participant. All quantities are expressed in degrees. *N* = 32. The pairs of panels in (A) and (B) represent the mean absolute change (left) and the standard deviation of change (right) for the two conditions. The error bars represent SEM over participants. Paired tests for significance between success and failure within (A) and between (B) tasks are reported. ^∗^ : *p <* 0.05, ^∗∗^ : *p <* 0.01, ^∗∗∗^ : *p <* 0.001

Moreover, the comparison between tasks (Fig.5B) shows significant differences due to the different feedback provided. The error task lead to a greater increase in variability and larger adjustments after failures compared to successes. This suggests that participants were more sensitive to failures in the Error task, driving more pronounced and consistent motor corrections.

### EEG Neural Metrics suggest mixture in Learning Mechanisms

During the experiments, we monitored the players’ brain activity to examine trends in Post-Movement Beta Rebound (PMBR) throughout their learning process. PMBR is defined as the peak in beta activity following movement cessation (see Fig. 6a) and is a well-established marker of sensorimotor cortex activation. It is associated with motor learning, and its trend is expected to depend on the underlying learning mechanism Tan et al. (2014, 2016); Torrecillos et al. (2015); Kranczioch et al. (2008); Haar and Faisal (2020).

**Fig 6.**
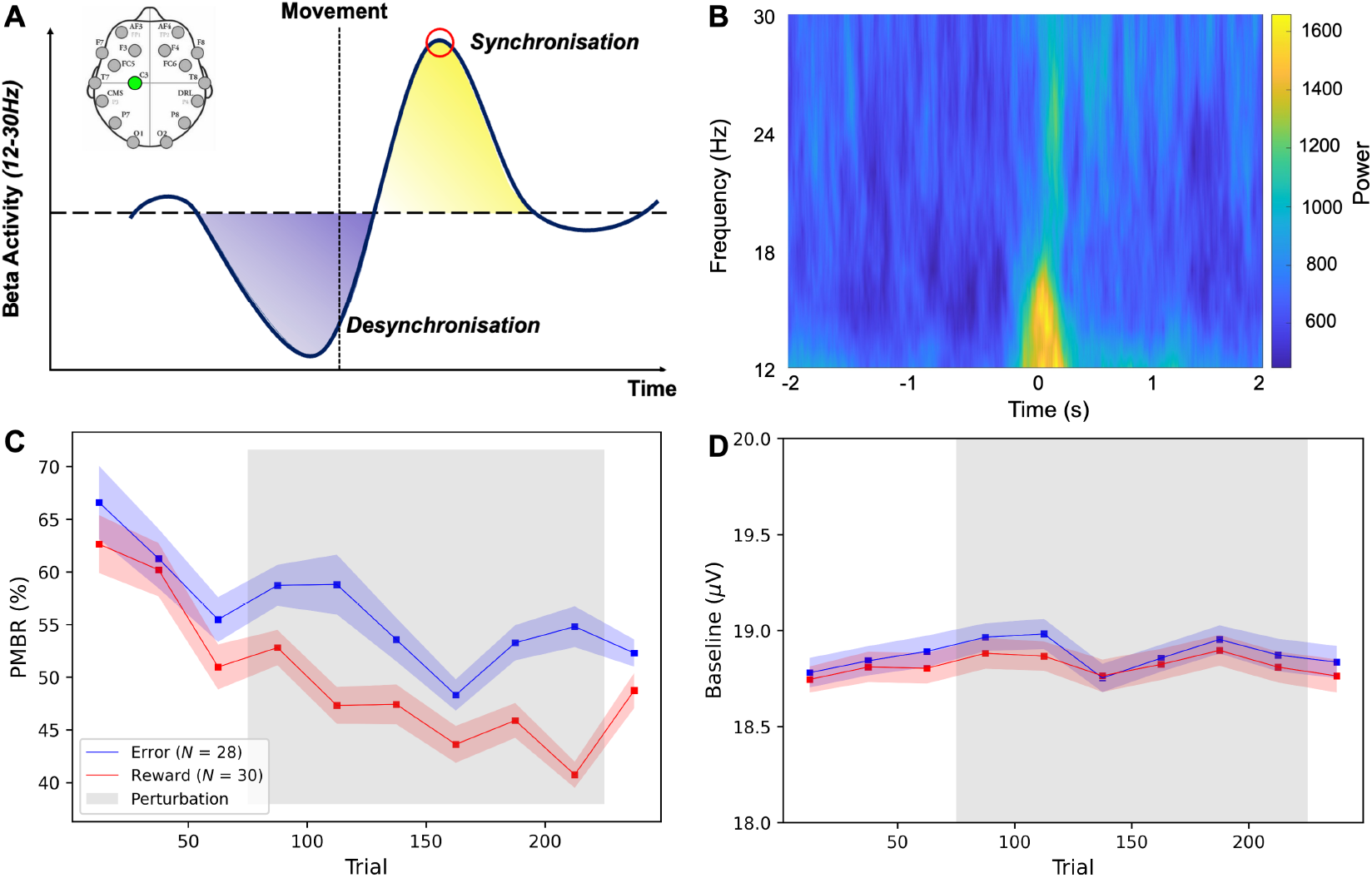
Post-Movement Beta Rebound. **(A)** Graphical representation of *β*-band *Event-Related Desynchronisation* (in blue) and *Event-Related Synchronisation* (in yellow). Circled in red is the *Post-Movement Beta Rebound* peak considered for the analysis. In the upper left angle, the *Emotiv* EPOC+ electrode map with C3 in green, the channel used for the PMBR derivation; **(B)** Time-frequency plot for an example session; **(C)** Average *Post-Movement Beta Rebound* ; **(D)** Average baseline power for the PMBR calculation. **(C-D)** All quantities are calculated within blocks of 25 trials over *error-based* condition (blue) and *reward-based* condition (red). The shaded areas represent the variability between the participants (standard error of the mean) and the grey area represents the presence of perturbation.

The increase in beta power was evident in the data (see fig. 6c) for all individual sessions. Both error and reward tasks exhibited similar PMBR trends during the baseline, with a consistent decrease over time. However, a distinct difference emerged in the PMBR trend during the rotation phase (fig.6b). The reward sessions showed a continuous decline (*b*_block_ = −1.955, *p <* 0.001, 95% CI: [-2.863, -1.047]), while participants in the error task did not exhibit a significant trend (*b*_block_ = −1.185, *p* = 0.09, 95% CI: [-2.534, 0.164]). During the washout, the average PMBR remained relatively consistent between conditions without significant differences (*F*_mode_(1, 56) = 0.69, *p* = 0.41). No significant baseline differences were identified between feedback types (*F*_mode_(1, 56) = 0.07, *p* = 0.79) or trends linked to the experimental phase (*F*_int_(9, 504) = 0.24, *p* = 0.99), ensuring that changes in PMBR were not driven by underlying shifts (fig. 6d).

When the data was split according to sessions, consistent with the behavioural measures, we observed the same trends as in the two-conditions split. The error task showed a non-significant trend, even though the magnitude differed between sessions (*b*_block|first_ = −2.019, *p* = 0.11, 95% CI: [-4.495, 0.458]; *b*_block|second_ = −0.462, *p* = 0.49, 95% CI: [-1.775, 0.851]). In contrast, the reward task displayed a significant decreasing trend over the learning process in both rounds (*b*_block|first_ = −2.405, *p <* 0.001, 95% CI: [-3.629 -1.180]; *b*_block|second_ = −1.504, *p* = 0.03, 95% CI: [-2.846, -0.163]).

## Discussion

In this study, we explored the impact of feedback manipulations on motor learning and the use of different learning mechanisms in a real-world task. By employing an Embodied Virtual Reality setup, we introduced a visuomotor perturbation and manipulated the feedback provided to participants during this real-world task. Our findings indicate that using either error-only or reward-only visual feedback was insufficient to induce learning solely based on error-based adaptation or reward-based skill acquisition, respectively. Both behavioural and neural data suggest that even when the reward feedback (ball pocketing) was removed in the error-only condition, participants still engaged in reward-based skill learning during this task. It seems that participants found sufficient rewards in observing what appeared to be the correct ball trajectory, similar to how professional basketball players celebrate successful shots before confirming if the ball goes into the net.

The results demonstrated a clear difference in how individuals adapted to the visuomotor perturbation when receiving error-only versus reward-only feedback. Participants adapted more rapidly to the cue ball rotation in the error feedback condition compared to the reward feedback condition, in accordance with previous research suggesting that adaptation to errors occurs faster than adaptation based on rewards Galea et al. (2015); Song et al. (2020). Furthermore, the difference in learning was even more pronounced between sessions. Participants who experienced the reward task first showed no learning, whereas those who had previously completed the error task were able to correct for most of the perturbation.

The definition of the reward zone played a key role here. The increase in received reward during the early rotation phase of the reward-only feedback condition did not encourage exploration and learning, as evidenced by the reduced inter-trial variability in those blocks. The dynamic reward zone, which decreased over time, made the task more challenging, leading to a decrease in perceived success and an increase in variability and exploration over the rotation blocks of the reward-only condition.

The dynamic reward provision was designed to guide participants on different paths without excessive or insufficient rewards, following established paradigms in lab-based motor learning tasks Therrien et al. (2016). However, due to this dynamic nature, participants did not experience an abrupt perturbation and instead noticed the changes slowly, which may have affected their perception of the rotated trajectory. As a result, in the reward-only feedback condition, the reduction in directional error exhibited a slow and linear learning curve. Similarly, the *real* success rate in the reward task showed a gradual, linear improvement mirroring the decline in directional errors. In contrast, the error-only feedback condition displayed a learning curve for the decrease in directional error that followed a double exponential pattern, consistent with typical adaptation tasks (e.g.,Taylor et al. (2014)) as well as performance in the realworld billiard task Haar et al. (2020). While feedback manipulations of error and reward are often thought to induce distinct learning mechanisms in lab-based tasks, our results suggest a more complex picture in this real-world setting.

Lag-1 autocorrelation [ACF(1)] of error has been proposed as a marker of expertise. For novices, it is expected to be significantly different from zero and decrease over time, as values closer to zero are associated with greater skill van Beers et al. (2013). The rationale is that as skills develop, individuals become less influenced by noise from their previous actions, and therefore the trials would be less correlated with the previous ones. In terms of the duality of learning mechanisms, ACF(1) is expected to decay during reward-based skill learning and remain unchanged during error-based adaptation.

During baseline, the ACF(1) did not differ from zero in both conditions. In the early learning phase, ACF increased and was significantly different from 0 in both conditions. Contrary to initial expectations, the error-only feedback condition exhibited a decrease in ACF towards zero during the perturbation phase, indicating reduced reliance on previous performance. This measure demonstrated an improvement in the reliability of the shot by the participants, suggesting that error-based adaptation was not the sole mechanism used for movement correction. The participants may have obtained reinforcement feedback from the expected outcome of the trajectory or through additional sensorimotor information, such as the proprioceptive feedback experienced when hitting the ball.

In contrast, the reward feedback condition displayed relatively consistent levels of ACF(1) over the rotation phase, implying a steady dependence on past results despite evolving task demands. This result can be attributed to the limited amount of learning in this condition or the continuous reliance on the binary reward information.

The decay in inter-trial variability, a measure traditionally associated with skill learning rather than adaptation, revealed intriguing differences between the error and reward conditions Müller and Sternad (2004); Cohen and Sternad (2009); Shmuelof et al. (2012); Huber et al. (2016). In the error condition, we observed a decline in inter-trial variability from the beginning to the end of the perturbation phase. Conversely, the reward condition displayed a more fluctuating and diverse pattern of inter-trial variability, without a clear decay trend across participants. This suggests a continuous exploration of different movement strategies, which aligns with the directional error and success rates in the reward-based feedback condition, indicating that participants were still learning by the end of the rotation phase.

Finally, the increased exploration after a failure in both tasks showed how reinforcement-based motor learning was used to learn the correct movements in both conditions.

Overall, these results converge to the idea that participants did not exclusively rely on the learning mechanism related to the specific feedback provided, especially in the error task, where there are clear signatures of reward-based skill learning.

To further validate the differences in learning measures between real-world and lab-based tasks, we also examined the neural activity of participants, specifically the Post-Movement Beta Rebound (PMBR) - the increase in power of neural beta oscillations (12-30Hz) at the end of movement.

PMBR is known to increase over learning in error-based adaptation tasks Tan et al. (2014, 2016); Torrecillos et al. (2015), whereas in skill-learning tasks, PMBR is expected to decrease (itself or its magnetic resonance spectroscopy correlate) over the learning Floyer-Lea et al. (2006); Kranczioch et al. (2008); Kolasinski et al. (2019). This divergence has been attributed to the projections of GABA activity for error- and reward-based learning from the cerebellum and basal ganglia, respectively Doyon et al. (2003); Doyon and Benali (2005).

Our results indicate that participants initially exhibited a high PMBR, which then decreased during the baseline sessions, as is typical of skill-learning tasks (e.g. Kranczioch et al. (2008)). This was expected, as participants were learning the EVR environment and the pool task. During the rotation phase, after the introduction of perturbation and partial feedback manipulation, the overall PMBR continued to decay for both feedback conditions. In the reward-only condition, there was a clear decay in PMBR, which is expected in a reward learning task without error-based adaptation. For the error-only condition, the PMBR trend was mixed, with a decay over the early learning phase suggesting the use of reward-based reinforcement skill learning. However, the PMBR increased over the later part of the rotation phase, where the errors likely became too small to reliably predict reward from the ball trajectory, leading to a shift towards implicit error-based adaptation. This implicit component is evident in the aftereffects observed during the washout phase.

Our findings suggest that participants engaged in a combined use of error- and reward-based learning in the real-world scenario, even when error or reward feedback was withdrawn. Both the behavioural and neural results indicate an interaction between these mechanisms, despite the attempts to isolate them through feedback manipulations. The richness of stimuli and experience in the real-world task appears to have limited the effects of these feedback manipulations. This is particularly evident in the error-only condition, where the participants likely had enough information from the cue ball trajectory to preserve their success and utilise reward-based reinforcement. Similarly, in the reward-only condition, while we withdrew visual sensory-error feedback, the participants still experienced full proprioceptive feedback, which could have guided some error-based adaptation.

While our study provides valuable insights into the interaction between error- and reward-based learning mechanisms in a real-world scenario, it is important to acknowledge certain limitations that may have influenced the findings. First, the sample size of participants could impact the generalisability of the results, and a larger and more diverse participant pool could offer a broader perspective on the dynamics of error- and reward-based learning in motor tasks. Second, the consideration of PMBR as a neural marker for the learning mechanisms may not have fully captured the complex neural processes associated with error- and reward-based learning. Future studies could explore a more comprehensive assessment of neural activity across different brain regions to provide a deeper understanding of the underlying mechanisms and identify patterns beyond PMBR. Third, the way the reward feedback was provided did not allow participants to fully correct the perturbation, due to the limited information available. Different methods to provide optimal reward information need further exploration, as the current reward zone method may be sufficient for lab-based tasks but could not be sufficiently informative in real-world scenarios. Finally, while the use of EVR provided a valuable context for studying error- and reward-based learning in real-world situations, it also introduced additional variables and complexities, such as the use of Virtual Reality, which may have influenced the observed dynamics.

The findings from this study challenge the notion that learning mechanisms can be isolated solely through feedback manipulation. The observed differences in ACF(1), variability decay, reward-dependent motor variability, and neural activity between error-only and reward-only tasks underscore the intricate nature of motor learning and adaptation in dynamic environments. While feedback manipulation shapes the learning process and influences motor performance, it is clear that participants do not rely exclusively on single learning mechanisms, even when receiving partial visual feedback. Instead, they engage in a holistic learning process that incorporates various environmental cues and internal computations.

In conclusion, this study provides valuable insights into the complexities of human motor learning and performance, emphasising the need for a nuanced understanding of the interplay between error-based and reward-based learning mechanisms. The findings have important implications for developing more effective learning interventions and rehabilitation strategies. Further research in this area holds the potential to advance our knowledge of motor learning processes and inform the design of interventions that better align with the inherent complexities of human motor adaptation.

## Declarations

### Ethics approval

All experimental procedures were approved by the Imperial College Research Ethics Committee and performed in accordance with the Declaration of Helsinki.

### Consent to participate

All subjects gave their informed consent before participating in the study.

### Availability of data and materials

Data will be made available on FigShare upon publication.

### Authors’ contributions

S.H. and A.A.F. conceived and designed the study; F.N., A.A.F. and S.H. developed the experimental setup; F.N. acquired the data; F.N., A.A.F. and S.H. analysed and interpreted the data; F.N. drafted the paper; SH and A.A.F. revised the paper.

